# CD200R1 Marks a Branch Point During Human Natural Killer Cell Development in Tonsils

**DOI:** 10.64898/2026.11.02.762613

**Authors:** Youssef Youssef, Ansel P. Nalin, Michael A. Ruesch, Matthew R. Lordo, Noah T. Koenigs, David Clever, Joel Cesar, Megan Broughton, Wing Chan, Anthony G. Mansour, Ekaterina Altynova, Xiaoli Zhang, James S. Blachly, Bradley W. Blaser, Christopher C. Oakes, Bethany L. Mundy-Bosse, Benjamin Krämer, Aharon G. Freud

**Affiliations:** Comprehensive Cancer Center, The Ohio State University, Columbus, OH, USA; Division of Hematology, Department of Internal Medicine, The Ohio State University, Columbus, OH, USA; Department of Radiation Oncology, The University of Texas MD Anderson Cancer Center, Houston, TX, USA; Medical Scientist Training Program, The Ohio State University, Columbus, OH, USA; Department of Pathology & Immunology, Washington University School of Medicine, St. Louis, MO, USA; Biomedical Sciences Graduate Program, The Ohio State University, Columbus, OH, USA; Division of Musculoskeletal Oncology, Department of Orthopaedic Surgery, Washington University School of Medicine, St. Louis, MO, USA; Department of Internal Medicine, Virginia Commonwealth University, Richmond, VA, USA; Department of Hematology and Hematopoietic Cell Transplantation, City of Hope National Medical Center, Los Angeles, CA, USA; Biostatistics Core, College of Nursing, College of Public Health, University of South Florida Health, Tampa, FL, USA; Department of Microbial Infection and Immunity, The Ohio State University, Columbus, OH, USA; Department of Internal Medicine I, University of Bonn, Bonn, Germany; German Center for Infection Research, Bonn, Germany; Department of Pathology, The Ohio State University, Columbus, OH, USA

**Keywords:** NK cell development, stage 4A NK cell, CD200R1, human ILC development

## Abstract

Human natural killer (NK) cells arise from common innate lymphoid cell progenitor and precursor populations that are selectively enriched within secondary lymphoid tissues and that complete their terminal differentiation through stages of progressively lineage-restricted downstream developmental intermediates. We identify two subsets of tonsil CD94/NKG2A^+^NKp80⁻ “stage 4A” NK cell development intermediates that show differential expression of the inhibitory receptor, CD200R1. Ex vivo single-cell transcriptomic, DNA methylation, phenotypic, and functional analyses demonstrate that CD200R1⁺ stage 4A cells represent an immature, oligopotent precursor population enriched for tissue-resident and cytokine-responsive programs. In contrast, CD200R1⁻ stage 4A cells show more restricted NK cell developmental potential concomitant with features of mature NK cells. These findings help to refine the human NK cell developmental pathway in tonsils and highlight CD200R1 as a useful marker delineating an uncommitted developmental state linking microenvironmental cues to NK cell maturation and plasticity.

## INTRODUCTION

Natural killer (NK) cells are founding members of the innate lymphoid cell (ILC) family that is composed of a diverse lineage of cytotoxic and immunomodulatory effector cells with non-redundant roles in host defense, tumor surveillance, tissue homeostasis, and pregnancy [1–5]. ILCs have been categorized into three major groups, each defined by differentially expressed surface receptors/antigens, transcription factors, and effector functions [6–8]. Group 1 ILCs include EOMES^+^T-BET^+^ NK cells, which can directly lyse virus-infected or malignant cells, as well as EOMES⁻T-BET^+^ ILC1s that are generally considered non-cytotoxic, immunoregulatory cells akin to CD4^+^ T-helper type 1 (Th1) cells. Group 2 ILCs include GATA3^+^ ILC2s that share several features with CD4^+^ Th2 cells; and group 3 ILCs include RORγt^+^ ILC3s that share features with CD4^+^ Th17 cells.

Previous studies support a model of human NK cell development whereby bone marrow-derived, multipotent, CD34^+^CD45RA^+^ hematopoietic progenitor cells (HPCs) migrate to peripheral tissues, including secondary lymphoid tissues (SLTs) such as tonsils and lymph nodes, to complete their differentiation and maturation [9]. At least six functionally distinct stages [aka NK cell developmental intermediates (NKDIs)] can be delineated among lineage (Lin) antigen-negative lymphocytes by the differential expression of surface markers including CD34, CD117, CD56, CD94, NKp80, CD16, and CD57 [10–12]. In tonsils, stage 1 (S1) NKDIs (CD34^+^CD117⁻) comprise the least differentiated NKDIs and retain the most developmental plasticity ex vivo (including potentials for T cell, dendritic cell, and other ILC differentiation) [11]. S2 NKDIs (CD34^+^ CD117^+^) are also multipotent yet include an IL-1R1^+^ subset of cells with ILC/NK-lineage restricted potential and referred to as common ILC progenitors [11, 13, 14]. S3 NKDIs comprise a heterogeneous pool of CD34⁻CD117^+^CD56^+/^⁻ cells that also show ILC/NK cell lineage restricted potentials and represent a major branch point from which divergent ILC/NK cell lineage trajectories directly emerge. Thus, S3 cells are frequently referred to as common ILC precursors (ILCPs) [10, 12, 13, 15, 16]. S4 NKDIs are more restricted still and are defined by the expression of the group 1 ILC/NK-associated markers, CD56 (bright) and CD94, in the absence of CD16 and CD57, which delineate terminally differentiated CD56^dim^ S5 and S6 NKDIs, respectively. Furthermore, S4 cells can be divided into two subsets based on the expression of the activating receptor, NKp80, which is specific to NK cells among ILCs: S4A (NKp80⁻) and S4B (NKp80^+^) [17]. Ex vivo, at least some S4A cells retain non-NK ILC potential, whereas S4B cells appear to represent the first stage of human NK cell lineage commitment and show mature NK cell effector functions ex vivo [12, 18].

The inhibitory cell surface transmembrane glycoprotein CD200 receptor 1 (CD200R1) is a key regulator of immune homeostasis and has been investigated as a target for immune-based therapy in several cancers [19–21]. CD200R1 interacts with its ligand (CD200), which is expressed on tumor cells and activated immune cells [22]. CD200R1 is normally highly expressed on various myeloid populations, and while it can be induced on some NK and T cell subsets following activation, in healthy blood and tissues, CD200R1 is typically restricted to non-NK ILCs [12, 16, 23]. Indeed, in both humans and mice, CD200R1 can be a useful marker distinguishing ILC1s (CD200R1^+^) from mature NK cells in adipose tissue, liver and other tissues [23–25]. However, the expression and functional roles of CD200R1 during earlier stages of ILC/NK development remain unexplored.

Here, we investigated the expression of CD200R1 among human NKDIs. Through single cell RNA sequencing (scRNA-seq) and flow cytometry studies of human tonsil NKDIs, we found that CD200R1 is highly (albeit nonuniformly) expressed during the S3-S4B developmental window. In general, we found that CD200R1 marked NKDIs with relative transcriptional, phenotypic and functional immaturity. Although we found no evidence to indicate that in vitro engagement of CD200R1 on S3 or S4A NKDIs with its ligand (CD200) influenced differentiation, we observed that CD200R1^+^ S4A cells were more developmentally plastic than their CD200R1^-^ counterparts, which showed many features of otherwise mature, terminally differentiated NK cells. Thus, CD200R1 expression refines the human NK cell developmental pathway and may serve as a useful marker of uncommitted intermediates with distinct differentiation and maturation capacities.

## RESULTS

### CD200R1 is expressed in intermediate tonsil NKDIs and marks a branch point in NK/ILC differentiation

To define the developmental timing and significance of CD200R1 expression during human ILC/NK cell development, we first analyzed a recently published scRNA-seq dataset generated from enriched tonsil NKDIs and ILCs [26]. Unsupervised UMAP projection of Lin⁻ cells revealed previously described NKDIs, ILC2s, and ILC3s as well as *ITGA1*^+^*ITGAE*^+^ tissue resident NK (trNK) cells (aka intraepithelial group 1 ILCs or ieILC1s) [8, 27, 28] identified based on expression patterns of known cell type-associated or -defining genes as well as by analysis of gene set modules generated from publicly available bulk RNAseq data (Figure 1A and Figure S1). As previously reported, we did not detect a distinct cluster of *EOMES*⁻*TBX21*^+^ ILC1s, likely due to their very low frequency in human tonsils [25]. Likewise, we did not detect a distinct cluster of S6 cells, which are also very rare in secondary lymphoid tissues [12, 29]. We next investigated the expression of *CD200R1* across the developmental landscape. UMAP visualization of gene expression revealed that *CD200R1* was expressed by most S3 and S4A NKDIs, ILC2s, and ILC3s, with little to no expression detected in early CD34^+^ HPCs (S1-2 cells), late-stage NKDIs, and ieILC1s/trNK cells (Figure 1B and Figure S1). This expression pattern positioned CD200R1^+^ cells at the previously described developmental branch point coinciding with the transition from multipotent S3 cells (ILCPs) toward lineage-committed S4B-5 NKDIs and trNK/ieILC1s (Figure 1C).

**Figure 1.**
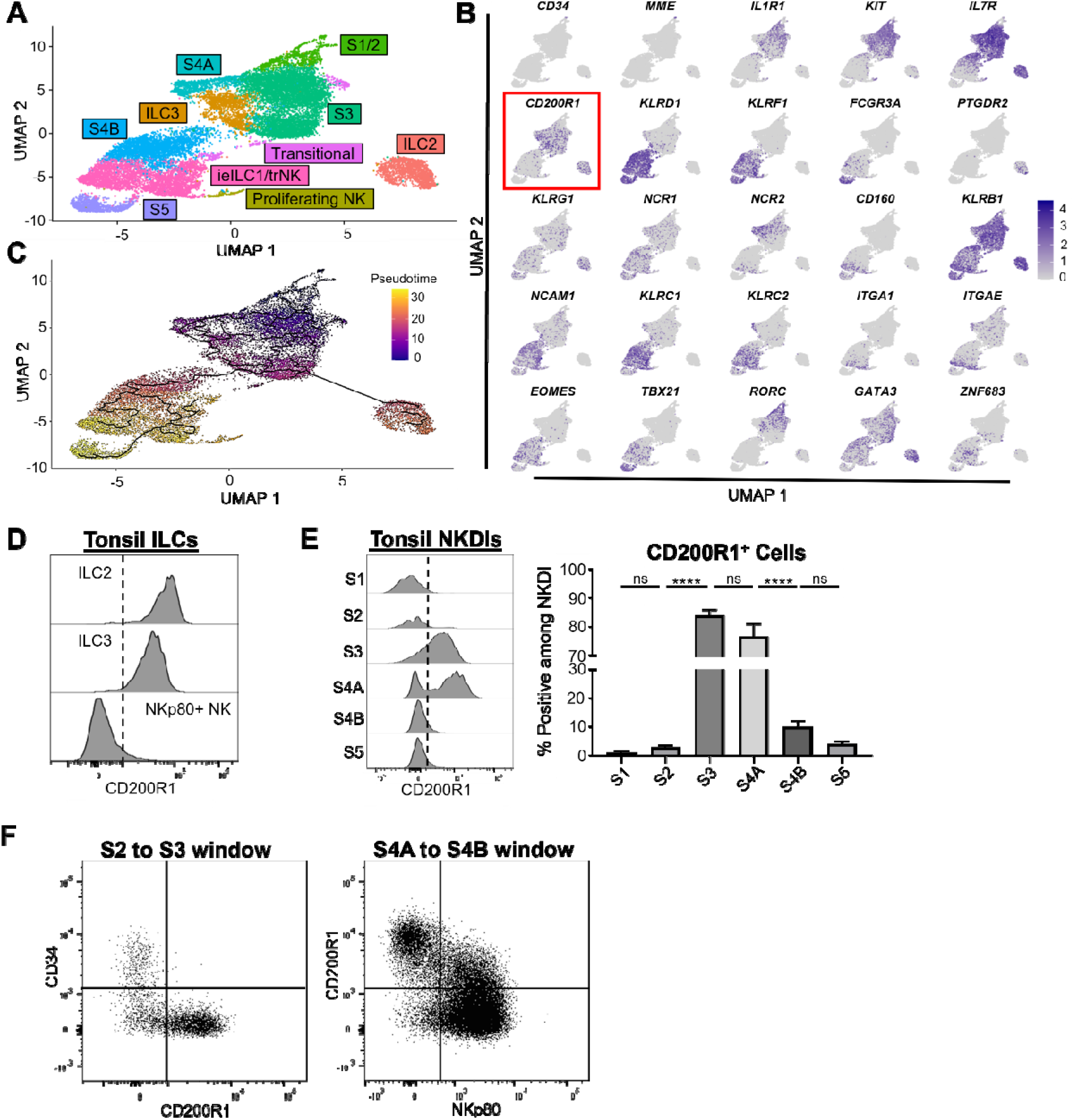
CD200R1 is expressed in intermediate tonsil NKDIs and marks a branch point in NK/ILC differentiation. (A) UMAP projection of previously published scRNA-seq data from tonsil NKDIs and ILCs [26], grouped by assigned cell type following Leiden-based clustering. Data are from two donors. (B) Feature plots of selected transcripts associated with NKDI/ILC identity and maturation. (C) Monocle 3 pseudotime trajectory analysis with CD34^+^ HPCs (S1 and S2 cells) designated as the root. (D) Representative histograms of CD200R1 surface expression by tonsil ILC2s, ILC3s, and NKp80⁺ NK cells. (E) Representative histograms of CD200R1 surface expression by the indicated ex vivo tonsillar NKDI subsets (left) and frequencies of CD200R1⁺ cells among these populations (right) (n=12). (F) Representative flow cytometry analyses of the S2-to-S3 developmental window (left) and the S4A-to-S4B developmental window (right). The dot plot on the left was gated on Lin⁻CD117⁺CD94⁻NKp44⁻CD294⁻KLRG1⁻NKp80⁻CD16⁻ cells. The dot plot on the right was gated on Lin⁻CD94⁺CD16⁻ cells. NKDIs and ILCs were defined as follows (all Lin^-^; Lin = CD3, CD4, CD5, CD14, CD19, CD20, CD123, FcεR1α, TCRαβ, TCRδγ): S1 (CD34⁺CD117⁻), S2 (CD34⁺CD117⁺), S3 (CD34⁻CD117⁺NKp44^-^CD294^-^KLRG1^-^CD94^-^NKp80^-^CD16^-^), S4A (CD94⁺NKp80⁻CD16⁻), S4B (CD94⁺NKp80⁺CD16⁻), S5 (CD94^+/-^NKp80^+^CD16^+^), ILC2 (CD94⁻NKp80^-^CD16^-^CD294⁺), ILC3 (CD94⁻NKp80^-^CD16^-^NKp44⁺). Data in (D) are representative of at least two independent experiments and are shown as the mean ± SEM. Statistical significance was determined using Student’s t test or one-way ANOVA followed by the Holm-Sidak method to correct for multiple comparisons. ns, not significant (p ≥ 0.05); ∗∗∗∗p ≤ 0.0001. See also Figure S1.

Ex vivo flow cytometry analyses of tonsil NKDIs and ILC subsets showed patterns of CD200R1 surface expression that matched the findings by RNAseq. As anticipated, tonsil ILC2s and ILC3s uniformly expressed CD200R1, with ILC2s showing higher mean fluorescence intensity expression (Figure 1D). Moreover, we observed that surface CD200R1 protein was expressed by most tonsil S3 and S4A cells, which showed the brightest expression among NKDIs. In contrast, CD200R1 was only dimly expressed by a minor subset of S4B cells, and it was not detected on S1, S2, or S5 cells (Figure 1E).

Given the heterogeneous expression of CD200R1 among S3-S4B NKDIs and to further define the dynamics of CD200R1 expression during ILC/NK cell development, we analyzed its expression in relation to surface markers that delineate key developmental transitions in tonsils. During the S2 (CD34^+^) to S3 (CD34⁻) developmental window (visualized by gating on total Lin⁻CD117^+^CD94⁻NKp44⁻CD294⁻KLRG1⁻NKp80⁻CD16⁻ cells), we observed that CD200R1 and CD34 exhibited mutually exclusive expression, indicating that CD200R1 is upregulated after, rather than during, the developmental transition from S2 to S3 (Figure 1F, left plot). Within the S4A (NKp80⁻) to S4B (NKp80^+^) developmental window (visualized by gated on total Lin⁻CD94^+^CD16⁻ cells), we noted a potential developmental trajectory stemming from the more abundant CD200R1^+^NKp80⁻ S4A subset toward the CD200R1⁻NKp80^+^ S4B population and progressing through a putative CD200R1^dim+^NKp80^+^ intermediate (Figure 1F, right plot). In contrast, the CD200R1⁻NKp80⁻ S4A fraction appeared to fall outside the above potential developmental pathway, raising the possibilities that CD200R1⁻ S4A cells either bypass a CD200R1^+^ trajectory or, at least in part, represent mature NK cells that may have downregulated NKp80 (and CD16) as can occur following activation [30, 31].

### CD200R1 expression in S3 NKDIs does not delineate distinct transcriptional identities or developmental potentials

Given the heterogeneity observed in the intermediate stage NKDIs (Figure 1E), we next sought to determine whether *CD200R1* expression delineated functionally distinct subpopulations within the S3 population. Unsupervised reclustering of the human tonsil scRNA-seq data shown in Figure 1 but limited to S3 cells revealed five transcriptionally distinct subclusters (Clusters 0-4), suggesting heterogeneity within this population (Figure 2A). However, when mapped onto these clusters, *CD200R1* expression was broadly distributed without specific enrichment in any one cluster (Figure 2B), indicating that *CD200R1*⁺ cells did not segregate into a distinct transcriptional subset of S3 cells.

**Figure 2.**
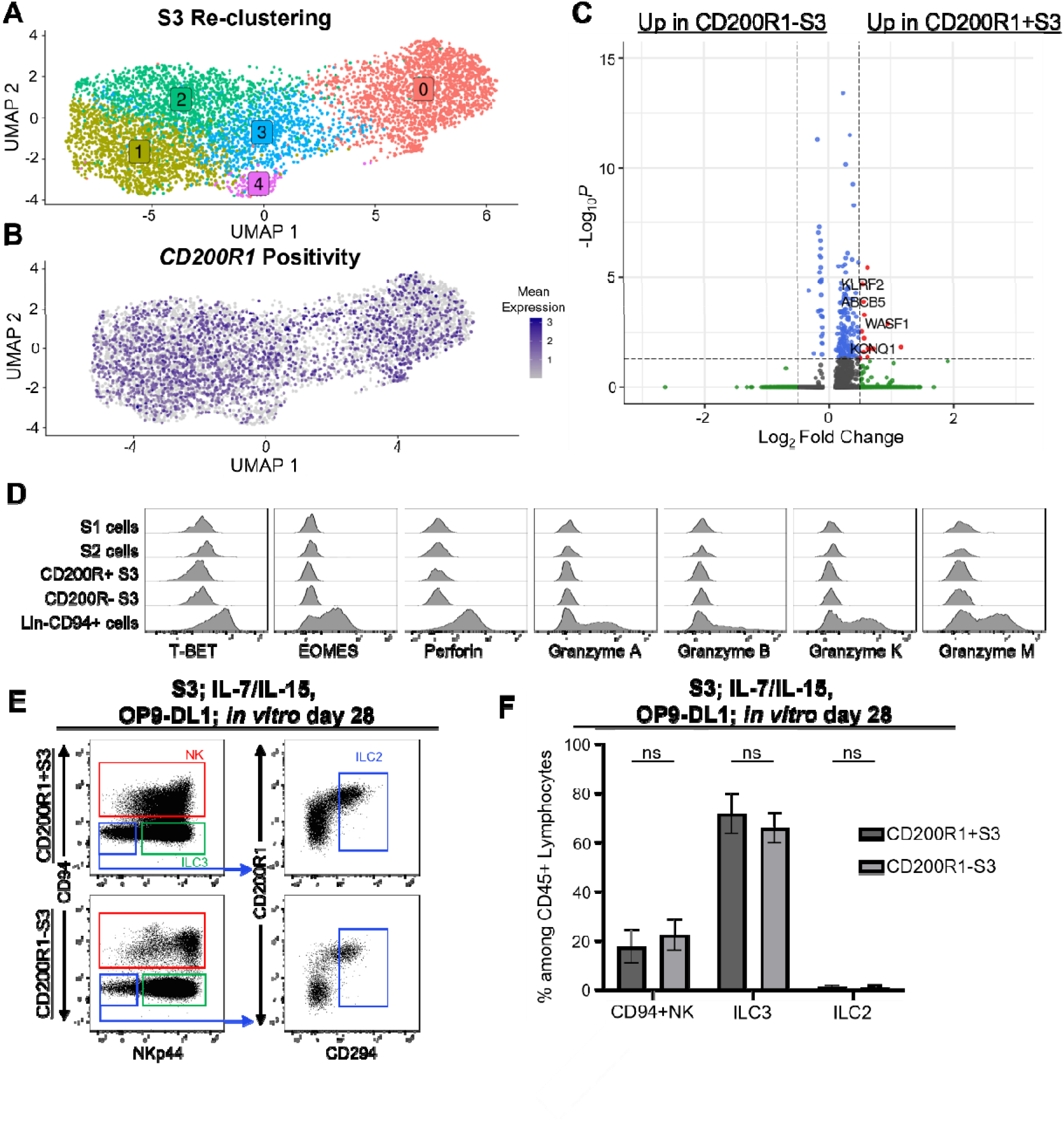
CD200R1 expression in S3 NKDIs does not delineate distinct transcriptional identities or developmental potentials. (A) UMAP projection of tonsillar S3 NKDIs following unsupervised reclustering, identifying five transcriptionally distinct clusters (0-4). (B) UMAP visualization of CD200R1 expression across the S3 clusters. (C) Volcano plot showing differentially expressed genes between *CD200R1*⁺ and *CD200R1*⁻ S3 NKDIs. (D) Representative flow cytometry histograms showing expression of the indicated markers by tonsil NKDIs. (E-F) Representative flow cytometry analyses (E) and frequencies (F) of NK cells, ILC3s, and ILC2s generated following 28-day culture of sorted CD200R1⁺ or CD200R1⁻ S3 NKDIs on OP9-DL1 stroma with IL-7 and IL-15 (n=6). Dot plots on the left were gated on live, Lin^-^ lymphocytes; dot plots on the right were gated on the CD94^-^NKp44^-^ fractions as indicated. NK cell, ILC3, and ILC2 subsets were identified as shown. Data are shown as the mean ± SEM. Statistical significance was determined using Student’s t test or one-way ANOVA followed by the Holm-Sidak method to correct for multiple comparisons. ns, not significant (p ≥ 0.05).

To determine if *CD200R1* expression correlated with NK cell lineage specification, we compared gene expression profiles between *CD200R1*⁺ and *CD200R1*⁻ S3 cells. Differential gene expression analysis revealed a small set of differentially expressed genes (DEGs) significantly enriched in the *CD200R1*^+^ group, including *ABCB5*, *KCNQ1, KLRF2*, and *WASF1* (Figure 2C). *ABCB5* was previously shown to reflect plasticity and stemness properties [32]; and *KLRF2*, which encodes for NKp65, was found to be selectively expressed on ILC3s but not on NK cells, ILC1s, or ILC2s [33]. Notably, we did not identify any canonical NK cell lineage-defining transcription factors or receptor genes, including *TBX21* (T-BET), *EOMES*, or *KLRD1* (CD94), that were differentially expressed. We also did not detect any appreciable acquisition of T-BET, EOMES, perforin, or granzyme proteins by either CD200R1⁺ or CD200R⁻ S3 subsets analyzed by ex vivo flow cytometry (Figure 2D). These findings suggest that *CD200R1* expression did not reflect any definitive NK cell lineage specification among S3 precursors, but its expression was potentially reflective of stemness and a bias towards ILC3 differentiation.

To test this possibility and determine if CD200R1 expression within S3 cells have distinct lineage differentiation potentials ex vivo, CD200R1⁺ and CD200R1⁻ S3 subsets were purified from fresh human tonsils and then cultured for four weeks with IL-7/IL-15 on OP9-DL1 stroma, a Notch ligand-expressing cell line that supports the differentiation of human ILCs and NK cells [11, 12, 34]. In these conditions, CD200R1⁺ and CD200R1⁻ S3 cells generated similar frequencies of NK cells (Lin^-^CD94^+^), ILC3s (Lin⁻CD94⁻CD294⁻NKp44⁺), and ILC2s (Lin⁻CD94⁻CD294⁺NKp44⁻) among total ILCs produced in culture (Figure 2E-2F).

### CD200R1^+^ S4A NKDIs are oligopotent and less terminally differentiated than CD200R1⁻ S4A counterparts

We next performed unsupervised clustering of scRNA-seq data from sorted total tonsil S4A NK cells. UMAP projection identified three transcriptionally distinct clusters within the S4A compartment, with clusters 1 and 2 demonstrating enrichment of *CD200R1* expression compared to cluster 3 (Figure 3A-3B). Clusters 1 and 2 were characterized by increased expression of genes associated with immature or ILC3-like programs, including *IL7R*, *NCR2* (NKp44), *KIT* (CD117), and *KLRB1* (CD161) [1, 10, 33]. Interestingly, cluster 1 showed relatively higher expression of *NCR2* and *KLRC1* (NKG2A), whereas cluster 2 showed relatively higher expression of *KIT* and *KLRB1*, highlighting heterogeneity within the CD200R1⁺ S4A compartment and potentially indicative of a spectrum of differentiation along a S3-to-NK developmental continuum. In contrast, cluster 3, which showed low *CD200R1* expression, exhibited higher expression of NK cell maturity and cytotoxicity associated genes, including *EOMES*, *TBX21* (T-BET), *PRF1* (perforin), *GZMB* (granzyme B), *NCR1* (NKp46), KIRs genes *(KIR2DL1, KIR2DL3, KIR2DL4, KIR3DL1,* and *KIR3DL2),* and *KLRC2* (encoding NKG2C), suggesting a more developmentally mature NK cell state [12]. Gene enrichment analysis further demonstrated that *CD200R1*⁺ S4A cells were enriched for S3 transcriptional programs, whereas *CD200R1*⁻ S4A cells were enriched for S5 gene signatures, further supporting their alignment with less and more mature states, respectively (Figure 3C).

**Figure 3.**
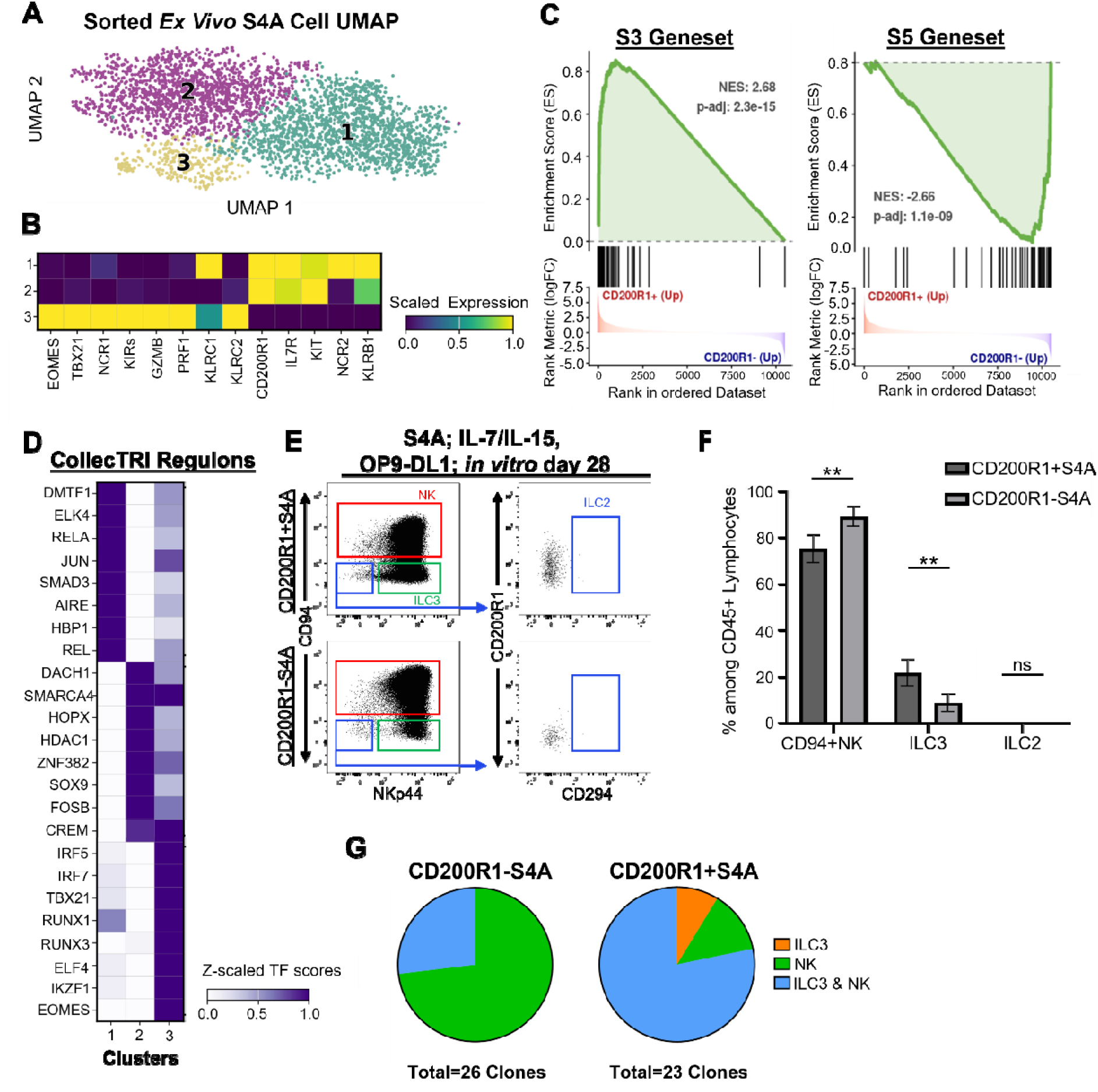
CD200R1^+^ S4A NKDIs are oligopotent and less terminally differentiated than CD200R1⁻ S4A counterparts. (A) UMAP projection of ex vivo FACS-purified tonsillar S4A NKDIs following unsupervised clustering. (B) Scaled expression of selected NKDI-associated genes among the three S4A clusters identified in (A). “KIRs” represents a combined gene expression score comprising *KIR2DL1*, *KIR2DL3*, *KIR2DL4*, *KIR3DL1*, and *KIR3DL2*. (C) Gene set enrichment analyses of DEGs between CD200R1⁺ and CD200R1⁻ S4A NKDIs using the S3 and S5 bulk RNAseq gene sets shown in Figure S1. (D) Scaled transcription factor activity scores for differentially active CollecTRI regulons among the three S4A clusters. (E-F) Representative flow cytometry analyses (E) and frequencies (F) of NK cells, ILC3s, and ILC2s generated following 28-day culture of sorted CD200R1⁺ or CD200R1⁻ S3 NKDIs on OP9-DL1 stroma with IL-7 and IL-15 (n=6). Dot plots on the left were gated on live, Lin^-^ lymphocytes; dot plots on the right were gated on the CD94^-^NKp44^-^ fractions as indicated. NK cell, ILC3, and ILC2 subsets were identified as shown. (G) Frequencies of single CD200R1⁺ or CD200R1⁻ S4A NKDI-derived clones producing ILC3s, NK cells, or both ILC3s and NK cells. A total of 23 CD200R1⁺ and 26 CD200R1⁻ S4A clones were analyzed from 4 donors in 2 separate experiments. Data in (F) are representative of at least two independent experiments and are shown as the mean ± SEM. Statistical significance was determined using Student’s t test or one-way ANOVA followed by the Holm-Sidak method to correct for multiple comparisons. ns, not significant (p ≥ 0.05); ∗∗p ≤ 0.01.

Transcription factor regulon analysis (CollecTRI) further revealed that the two *CD200R1*⁺ S4A clusters were not transcriptionally identical. Cluster 1 showed enrichment of *SMAD3-*, AP-1- and NF-κB-associated regulons, including those for *JUN*, *RELA*, and *ELK4* (Figure 3D) [35–37]. Cluster 2 showed enrichment of transcriptional regulators associated with chromatin remodeling, developmental cell-fate specification, tissue adaptation, and cellular plasticity (*DACH1, SMARCA4, HOPX, HDAC1, SOX9, FOSB,* and *CREM*) (Figure 3D) [38–43]. In contrast, cluster 3 cells (low for *CD200R1* expression) were enriched for canonical NK cell maturation-associated regulons, including those for *TBX21, EOMES, and RUNX* family members, again consistent with a more developmentally mature and cytotoxic NK cell program [44, 45].

Having observed that the absence of CD200R1 expression among S4A cells was associated with a more mature NK cell transcriptional profile, we sought to determine if CD200R1^+^ and CD200R1⁻ ILC/NK cell developmental intermediates have distinct lineage differentiation potentials in vitro. To test this, CD200R1⁺ and CD200R1⁻ subsets of S4A cells were purified from fresh human tonsils and then cultured for four weeks with IL-7/IL-15 on OP9-DL1 stroma. Unlike CD200R1^+^ and CD200R1⁻ S3 cells (Figure 2E-F), neither CD200R1⁺ nor CD200R1⁻ S4A cells produced ILC2s, consistent with prior findings [17, 18]. However, in bulk cultures, CD200R1⁻ S4A cells generated more NK cells and fewer ILC3s than CD200R1⁺ S4A cells (Figure 3E-3F). To confirm the bulk culture findings and determine the precursor frequencies within the S4A fractions, we performed single-cell clonal assays starting with tonsil CD200R1^+^ and CD200R1⁻ S4A subsets. Individually sorted CD200R1⁺ and CD200R1⁻ S4A cells were cultured with IL-7, IL-15, and OP9-DL1 stroma for 4 weeks. Most clones derived from CD200R1⁺ cells produced mixed populations of ILC3s and NK cells, and few clones contained exclusively ILC3s (Figure 3G). In contrast, only a minority of clones derived from CD200R1⁻ S4A cells contained ILC3s; and none of the CD200R1⁻ clones exclusively contained ILC3s (Figure 3G). Together, these findings demonstrate that most CD200R1⁺ S4A cells retain oligopotent ILC3/NK-developmental potential, whereas most CD200R1⁻ S4A cells are more restricted toward the NK cell lineage, supporting CD200R1 as a marker of a less terminally differentiated S4A NKDI state.

### CD200R1⁻ S4A cells show mature DNA methylation, phenotypic, and functional features

Given our transcriptional and in vitro culture findings comparing CD200R1^+^ and CD200R1⁻ fractions within the tonsil S4A compartment, we investigated other parameters of maturation. DNA methylation can serve as a stable indicator of maturational state [46, 47]. We purified tonsil S4A subsets and generated whole genome DNA methylation sequencing data as we had previously performed with tonsil S1-S5 NKDIs [46, 48]. Ex vivo profiling showed significantly higher calculated DNA methylation-based NK cell development scores for CD200R1⁻ S4A cells compared to CD200R1^+^ S4A cells, with the scores of CD200R1⁻ S4A cells closely approximating the scores of tonsil S4B cells (Figure 4A). Consistent with these findings, unsupervised clustering of differentially methylated loci, visualized as a heatmap, showed that CD200R1⁻ S4A cells shared features with more mature S4B and S5 NK cells, including regions associated with *CD160, FASLG, IL18RAP, NKG7,* and *TBX21* (Figure 4B). In contrast, CD200R1⁺ S4A cells showed relative enrichment of methylation patterns associated with cytokine-responsive and less mature S3 NKDIs, including regions associated with *IL1R1, IL4R, IL7R,* and *SMAD3* [7, 8, 12, 23, 49].

**Figure 4.**
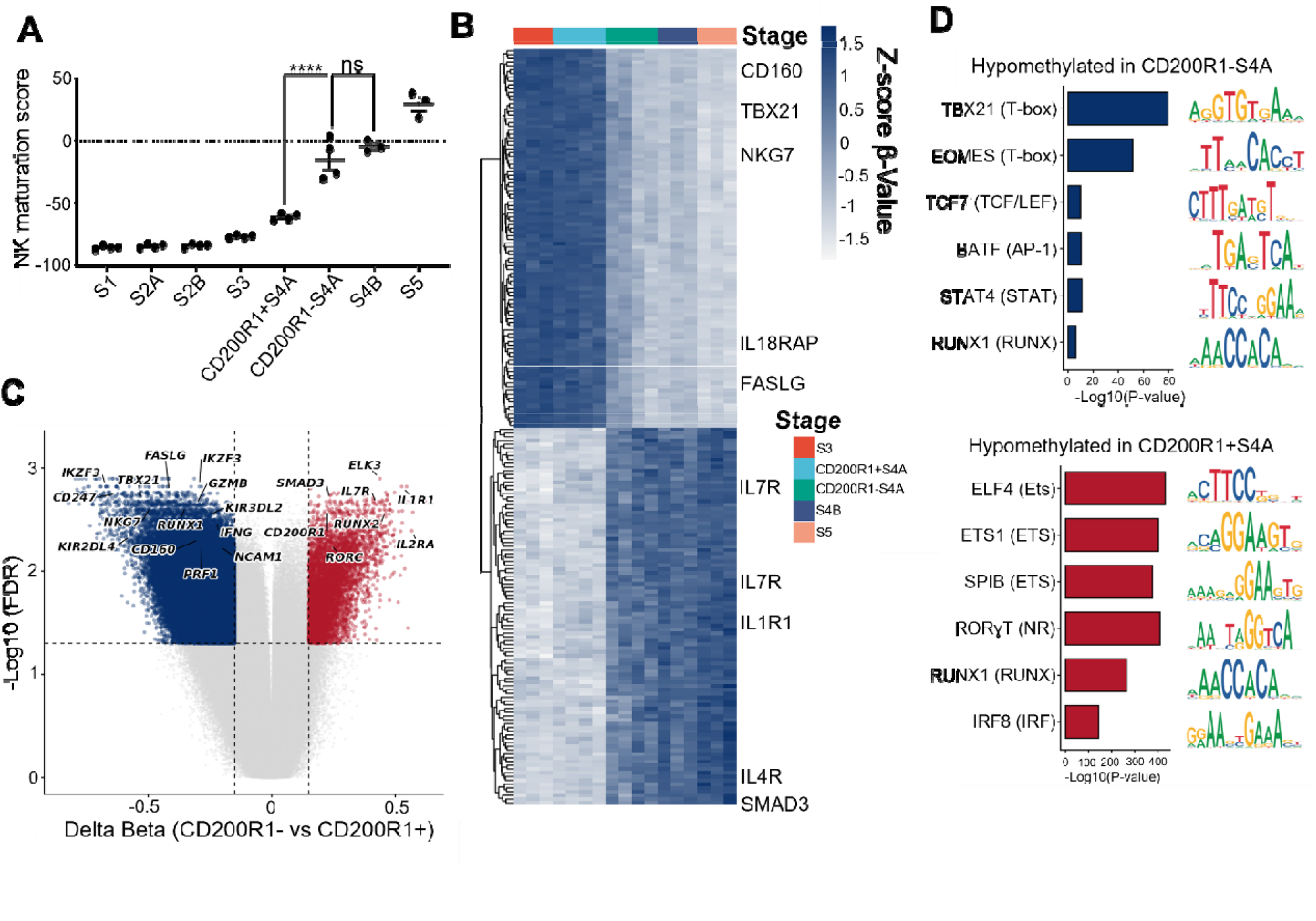
CD200R1⁻ S4A cells show mature DNA methylation features. (A) DNA methylation-based NK cell maturation scores among tonsil-derived NKDIs. Methylation scores are shown in arbitrary units and represent the first principal component (PC1) of a principal component analysis (PCA) using the 5,000 most-variable CpGs among NKDIs as previously described [46]. (B) Heatmap of the top probes contributing in each direction to PC1 of the PCA described in (A). β-values are displayed as Z-scores ranging from −1.5 to 1.5, with higher values indicating relative hypermethylation and lower values indicating relative hypomethylation. Some relevant genes associated with these probes are labeled. (C) Volcano plot of differentially methylated probes comparing CD200R1⁻ and CD200R1⁺ S4A cells. Significance was defined as an absolute Δβ >0.15 and FDR <0.05, as indicated by the dashed lines. Relevant genes associated with differentially methylated probes are labeled. (D) Motif enrichment analysis of regions hypomethylated in CD200R1^-^ (top) and CD200R1^+^ (bottom) S4A NKDIs. Motif enrichment was performed using HOMER de novo motif analysis, and the top enriched motifs from each analysis are shown. Statistical significance of motif enrichment is displayed as −log₁₀(P value), with the corresponding motif shown to the right of each bar. Transcription factor families are indicated in parentheses. ∗∗∗∗p ≤ 0.0001

Differential DNA methylation analysis, displayed as a volcano plot, further indicated broad epigenetic divergence between CD200R1^+^ and CD200R1⁻ S4A cells (Figure 4C). Using a delta β cutoff >0.15 and FDR <0.05, regions hypomethylated in CD200R1⁻ S4A cells were enriched near genes involved in NK cell cytotoxicity and terminal maturation, including genes encoding AIOLOS (*IKZF3*), KIRs, perforin (*PRF1*), and T-BET (*TBX21*) [12, 17, 46, 50]. In contrast, regions hypomethylated in CD200R1⁺ S4A cells were enriched near genes linked to immaturity, ILC3s, and/or tissue residency (*IL1R1*, *IL2RA*, *RORC*, *RUNX2, SMAD3*) [6, 51–53] (Figure 4C). Motif enrichment analysis supported this distinction, with *TBX21* and *EOMES* motifs enriched among regions hypomethylated in CD200R1⁻ S4A cells, while *RORγT, RUNX1, and SPIC* motifs were enriched among regions hypomethylated in CD200R1⁺ S4A cells as identified by de novo motif analysis (HOMER) (Figure 4D).

To investigate possible differences at the protein level, we performed additional ex vivo flow cytometry analyses of NK cell maturation-associated markers among tonsil NKDIs (Figure 5A). Interestingly, CD200R1⁻ S4A cells exhibited significantly less NKG2A surface expression compared to CD200R1^+^ S4A cells. In contrast, compared to CD200R1⁺ S4A cells, CD200R1⁻ S4A cells expressed significantly more NKG2C, KIRs (detected by pan-KIR2D and pan-KIR3DL1/L2 antibodies), EOMES, T-BET, perforin, and granzymes. Notably, the frequencies of NKG2C⁺, KIR⁺, and T-BET⁺ cells within the CD200R1⁻ S4A population were also significantly higher than those observed in CD200R1⁺ S4B cells, and for several of the markers the frequencies of the CD200R1⁻ S4A population were comparable to or approached those of CD200R1⁻ S4B and even S5 NK cells (e.g. NKG2A, NKG2C, KIR, and granzyme B).

**Figure 5.**
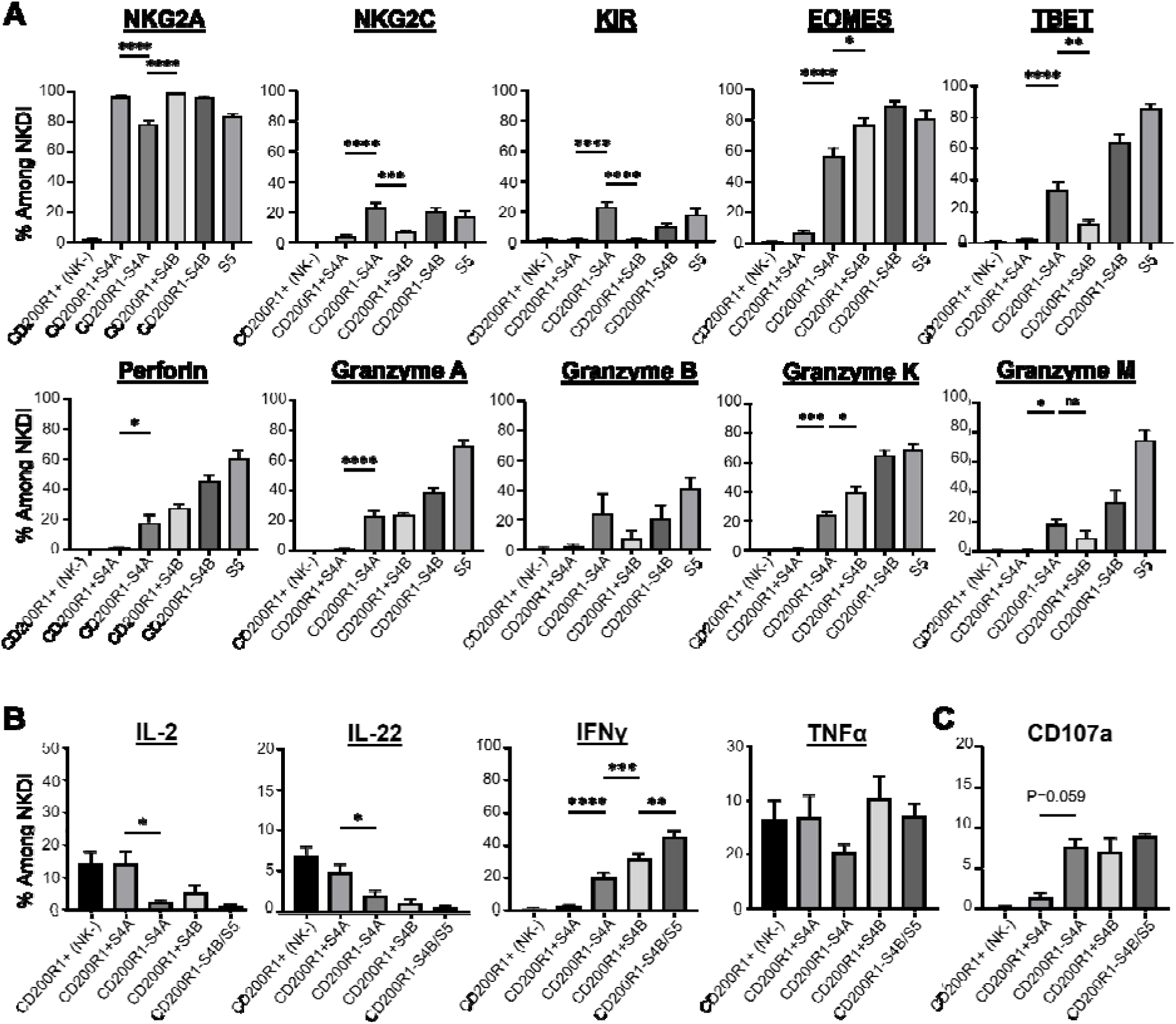
CD200R1^-^ S4A cells show mature phenotypic and functional features. (A) Frequencies of the indicated tonsil-derived cell populations expressing the indicated markers as measured by ex vivo flow cytometry. NKG2A-KIR, n=16; EOMES-granzyme A and granzyme K, n=6; granzyme B and granzyme M, n=4. CD200R1⁺ (NK^-^) cells were defined as Lin⁻CD94⁻NKp80⁻CD200R1⁺ cells, thus comprising a combination of S3 cells, ILC3s, and ILC2s. KIR expression was assessed using pan-KIR2D and pan-KIR3DL1/L2 antibodies. (B-C) Frequencies of the indicated tonsillar NKDI populations expressing the indicated cytokines as assessed by flow cytometry following negative selection enrichment of ILCs and NKDIs, overnight culture in cytokine-free medium, and four-hour stimulation with either IL-2/PMA/ionomycin (B) or IL-2 and K562 cells (C). IL-2, n=5; IL-22, n=9; IFNγ, n=17; TNFα, n=8; CD107a, n=4. S4B/5 cells were grouped together in these analyses and defined as total Lin⁻NKp80^+^ cells accounting for the downregulation of CD16 that occurs following stimulation. Data are representative of at least two independent experiments and are shown as the mean ± SEM. Statistical significance was determined using Student’s t test or one-way ANOVA followed by the Holm-Sidak method to correct for multiple comparisons. ns, not significant; ∗p ≤ 0.05, ∗∗p ≤ 0.01, ∗∗∗p ≤ 0.001, ∗∗∗∗p ≤ 0.0001.

We also assessed functional attributes of the S4A subsets and compared these to other NKDIs from fresh tonsil samples. Total ILCs and NKDIs were enriched by negative selection, cultured overnight in media without cytokines, and then stimulated for four hours with either IL-2/PMA/ionomycin to assess for cytokine production (Figure 5B) or with MHC class I negative K562 cells to assess for target cell induced degranulation (Figure 5C). Under these conditions, CD200R1⁻ S4A cells produced significantly less IL-22 and IL-2 and significantly more IFNγ than CD200R1⁺ S4A cells. However, CD200R1⁻ S4A cells produced less than NKp80⁺ NK cells (Figure 5B) (note: due to downregulation of CD16 that often occurs following activation [30, 31, 54], we could not reliably distinguish S4B from S5 cells). TNFα production was not significantly different across subsets (Figure 5B). In response to co-culture with K562 target cells, CD200R1⁻ S4A cells showed a near-significant trend toward increased degranulation compared with CD200R1⁺ S4A cells and like the amounts of NKp80⁺ NK cells (Figure 5C). Collectively, these DNA methylation, phenotypic, and functional features indicate that despite the lack of detectable NKp80 and CD16 surface expression, the tonsil CD200R1⁻ S4A population shows many characteristics of mature NK cells. In contrast, the CD200R1^+^ S4A population shows features of developmental immaturity and consistent with a precursor phenotype.

### Inferred niche signaling associates CD200R1⁺ S4A cells with epithelial, stromal, and myeloid-derived cues

Considering the distinct phenotypes of the CD200R1^+^ and CD200R1^-^ S4A subsets, we used the pediatric tonsil scRNA-seq dataset described in Figure 3 in conjunction with two publicly available scRNA-seq tonsil datasets [55, 56]. to perform a ligand-receptor interaction analysis comparing CD200R1⁺ and CD200R1⁻ S4A cells. Differential ligand analysis identified distinct sets of niche-associated ligands predicted to interact with CD200R1⁺ (right) and CD200R1⁻ (left) S4A cells within the tonsillar microenvironment (Figure 6A). CD200R1⁺ S4A cells were associated with ligands involved in adhesion (LGALS1, ITGB2) [57, 58], and immune regulation including CD200 (the CD200R1 ligand) as well as CLEC2D (a CD161 ligand) [59], BAG6 (an NKp30 ligand) [60], CD48 (a 2B4 ligand) [61], and MIF (through CXCR4) [62]. In contrast, CD200R1⁻ S4A cells were associated with a different set of ligands, including HLA-B (can engage inhibitory KIRs, including KIR3DL1 through HLA-Bw4) [63], B2M (a structural component of MHC class I complexes) [64], ICAM3 (a CD11d/CD18 ligand) [65], and HMGB1 (involved in NK-dendritic cell cross-talk) [66] (Figure 6A). Gene ontology (GO) biological process analysis of differentially interacting ligands and receptors also revealed distinct functional programs between the two niches. Ligand-associated pathways (Figure S2A) included processes such as chemokine production, inflammatory response, cytokine stimulation, cell-cell adhesion, and ERK1/2 signaling. Receptor-associated pathways (Figure S2B) indicated that CD200R1⁺ S4A cells are predicted to receive signals linked to immune response, lymphocyte activation and proliferation, integrin-mediated signaling, adhesion, cytokine production, and regulation of NK/leukocyte-mediated cytotoxicity.

**Figure 6.**
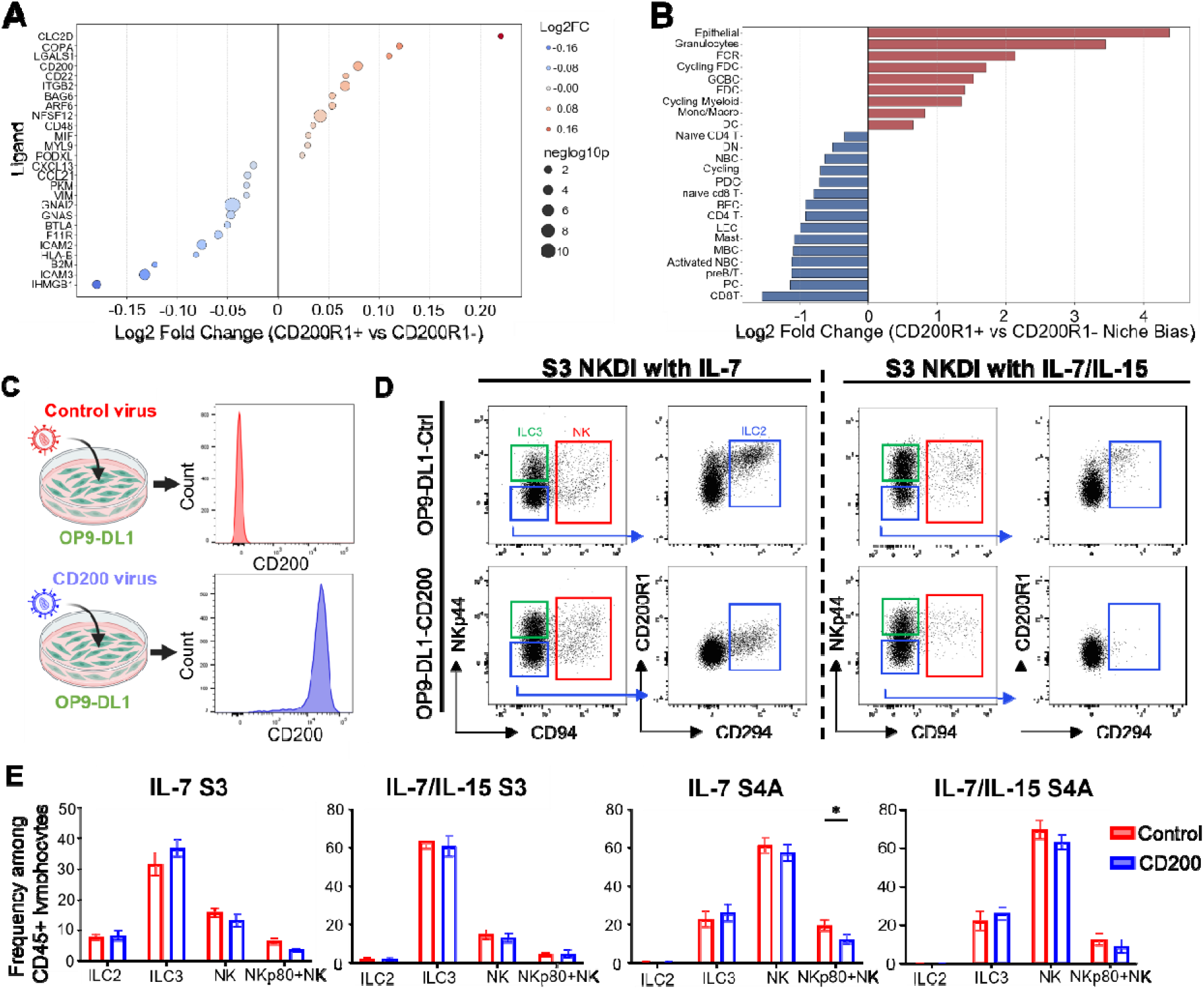
Inferred niche signaling associates CD200R1⁺ S4A cells with epithelial, stromal, and myeloid-derived cues. (A) Differential ligand interaction analysis comparing CD200R1⁺ and CD200R1⁻ S4A NKDIs. Positive log₂ fold change values indicate ligands preferentially associated with CD200R1⁺ S4A cells, whereas negative values indicate ligands preferentially associated with CD200R1⁻ S4A cells. Point size represents −log₁₀(P value). (B) Predicted cellular sources contributing to differential niche signaling received by CD200R1⁺ and CD200R1⁻ S4A NKDIs. Bars represent log₂ fold change in niche bias between CD200R1⁺ and CD200R1⁻ S4A cells. Positive values indicate signaling preferentially directed toward CD200R1⁺ S4A cells, whereas negative values indicate signaling preferentially directed toward CD200R1⁻ S4A cells. (C) Schematic depicting the generation of control and human CD200-expressing OP9-DL1 stromal cells, with representative flow cytometry histograms confirming CD200 surface expression on the OP9-DL1 cells. (D-E) Representative flow cytometry analyses (D) and frequencies (E) of NK cells, ILC3s, and ILC2s generated from total S3 and S4A NKDIs cultured on control or CD200-expressing OP9-DL1 stroma in the presence of IL-7 or IL-7/IL-15 for 28 days (n=9). Dot plots on the left were gated on live, Lin^-^ lymphocytes; dot plots on the right were gated on the CD94^-^NKp44^-^ fractions as indicated. NK cell (red boxes), ILC3 (green boxes), and ILC2 (blue boxes) subsets were identified as shown. Data are shown as the mean ± SEM. Statistical significance was determined using Student’s t test or one-way ANOVA followed by the Holm-Sidak method to correct for multiple comparisons. ∗p ≤ 0.05. See also Figure S2.

We next assessed the predicted cellular sources contributing to these niche interactions. Microenvironmental cell types, including epithelial cells, granulocytes, fibroblastic reticular cells (FRCs), follicular dendritic cells (FDCs), myeloid populations, monocytes/macrophages, and dendritic cells, were predicted to preferentially interact with CD200R1⁺ S4A cells (Figure 6B, right/red). In contrast, lymphoid populations, including CD8 T cells, CD4 T cells, plasma cells, B-lineage cells, and NK cells, were predicted to preferentially interact with CD200R1⁻ S4A cells (Figure 6B, left/blue).

Given these predicted ligand-receptor interactions, and considering the results of the culture studies described earlier (Figures 2 and 3), we investigated one relevant ligand-receptor pair: CD200-CD200R1. To determine whether CD200R1 signaling impacts differentiation, we cultured freshly isolated total S3 and total S4A cells from human tonsils (irrespective of CD200R1 expression to avoid labeling the cells with CD200R1 antibody during the sorts) with either IL-7 or IL-7/IL-15 and with OP9-DL1 stroma engineered to stably overexpress the CD200 ligand. The surface expression of the ligand was validated by flow cytometry (Figure 6C); and following co-culture of S3 cells with OP9-DL1-CD200 stroma, the resultant ILC2s showed reduced surface expression of CD200R1, consistent with receptor-ligand engagement and subsequent downregulation/internalization (Figure 6D). Comparing percentages of ILC subsets generated in the presence of control OP9-DL1 stroma versus OP9-DL1-CD200 stroma, we observed no significant differences in terms of ILC2, ILC3, and total NK cell production. We did note mildly decreased proportions of NKp80^+^ NK cells (i.e. S4B cells) produced in the presence of CD200, and the difference reached statistical significance in the IL-7 condition starting with S4A cells (Figure 6E).

Together, these data indicate that CD200R1⁺ and CD200R1⁻ S4A subsets occupy distinct inferred signaling niches within tonsillar SLT, with CD200R1⁺ cells preferentially associated with epithelial, stromal, granulocytic, and myeloid-derived interactions, whereas CD200R1⁻ cells are more strongly associated with lymphoid-derived interactions and HLA class I-KIR signaling, consistent with their increased KIR expression and more mature NK cell phenotype.

## DISCUSSION

NK cell development involves multiple concurrent processes, including differentiation, maturation, and diversification, which act in concert and are influenced by both cell intrinsic developmental plasticity and cell extrinsic signals stemming from the microenvironment [67]. Considering the latter, accumulating evidence suggests that at least some human NK cells arise from bone marrow-derived HPCs that circulate in the blood and then seed and terminally differentiate in peripheral tissues, which may confer tissue-specific phenotypes to locally derived NK cells [68–72]. Although not necessarily representative of NK cell development in all tissues, several prior studies focusing on NKDIs found within lymph nodes and tonsils have provided a working model of the stages of human NK cell development [9, 17, 73]. Notably, we and others have demonstrated that prior to S4B, intermediates along this NK-centric developmental continuum can also differentiate into non-NK ILCs [10–15, 18, 34, 74]. Indeed, investigation is ongoing to refine the closely connected NK cell and ILC developmental pathways, allowing for mechanistic elucidation and ultimately providing insights for clinical applications such as high-fidelity therapeutic targeting and adoptive cell therapy [48, 75–77].

In this context, here we evaluated the expression of the inhibitor receptor, CD200R1, among previously identified human tonsil NKDIs. Our data demonstrate that CD200R1 expression was primarily restricted to intermediate stage NKDIs, with abundant expression observed at S3 and S4A and with low expression detected on a minority of S4B cells. Because CD200R1 expression was heterogeneous at these stages and because prior studies in mice and humans had suggested that CD200R1 is a distinguishing marker of non-NK ILCs [24, 25, 78, 79], we investigated whether CD200R1 expression identifies developmental intermediates with distinct differentiation potentials. Within the S3 compartment, we found no clear evidence to suggest that CD200R1 expression was associated with restriction toward one or more specific ILC lineages. However, our studies did uncover previously unappreciated heterogeneity within the S4A population. In particular, the CD200R1^+^ S4A fraction showed DNA methylation, transcriptional, immunophenotypic, and functional features indicative of uncommitted NK cell precursors whose overall phenotype could position them along a developmental continuum between CD200R1^+^ S3 and CD200R1^dim+^ S4B cells and ultimately leading towards CD200R1^-^ S4B and S5 cells. In contrast, most cells within the minor CD200R1^-^ S4A population showed ex vivo features characteristic of a mature NK cell phenotype. Indeed, despite the traditional “S4A” designation of these cells based on their expression of CD94 in the absence of detectable NKp80 and CD16 expression [17], the CD200R1^-^ S4A population consisted mostly of cells with more mature transcriptional, DNA methylation-based, and functional features; and most CD200R1^-^ S4A cells showed NK cell lineage restriction in single cell clonal assays. Moreover, given their high amounts of NKG2C and KIRs along with relatively low amounts of NKG2A at the population level, most CD200R1^-^ S4A cells showed a phenotype that – based on our contemporary understanding of how NKG2A, NKG2C, and KIRs are regulated – would otherwise position them out-of-sync with the S3 to S4B continuum and would instead place them towards the end of the developmental spectrum [50, 72, 80]. Nonetheless, we observed that at least some CD200R1^-^ S4A cells showed developmental plasticity and ILC3 precursor potential, like most of the CD200R1^+^ S4A population. Thus, we favor that while most developing NK cells (and ILC2s and ILC3s) in tonsils proceed through a CD200R1^+^ phase of development, rare developing NK cells may bypass a CD200R1^+^ phase or may downregulate CD200R1 prior to acquiring detectable expression of NKp80 at S4B.

Beyond their intrinsic differences in maturation-associated gene expression and in vitro developmental potentials, our ligand-receptor analyses suggest that CD200R1⁺ and CD200R1⁻ S4A cells occupy distinct communication niches within the tonsil microenvironment. Notably, recent spatial profiling studies suggest that within human tonsils NKDIs show different locations and cell-cell interactions according to developmental stage [81]. Considering this prior work, our analyses predict that CD200R1⁺ cells preferentially interact with epithelial, stromal, follicular dendritic, granulocytic, and myeloid populations through signaling pathways associated with cytokine responsiveness, cell adhesion, developmental regulation, and tissue organization. In contrast, our analyses predict that CD200R1⁻ cells may preferentially engage other lymphoid populations through MHC class I-associated ligand-receptor interactions, including HLA-B/B2M complexes that may engage inhibitory NK cell receptors such as KIRs, together with signaling pathways associated with immune activation and mature NK cell function. Interestingly, these niche-specific interactions closely aligned with transcriptional programs identified by regulon analysis, whereby CD200R1⁺ cells were enriched for SMAD3-, AP-1-, and tissue-associated regulatory networks, suggesting that local microenvironmental cues contribute to establishing or maintaining their developmental states.

Historically, cytokine signaling pathways have been implicated as primary regulators of ILC differentiation and maturation [82–84]. Nonetheless, as NK cells and other ILCs develop, they express a multitude of additional surface receptors, cell adhesion molecules, and intracellular ligand-activated transcription factors that may influence developmental plasticity and fate determination. For example, we and others previously demonstrated that interactions with stromal cells and intracellular signaling pathways stemming from Notch receptors and the aryl hydrocarbon receptor regulate human ILC differentiation and maturation [12, 17, 18, 85–88]. To date, CD200R1 has primarily been studied as an inhibitory receptor that modulates immune-cell effector functions, including mature NK cell cytotoxicity [19, 21, 22, 89–91]. However, to the best of our knowledge, the potential role of CD200R1 during human NK cell development has not previously been investigated. In our studies, engagement of CD200R1 by CD200 had only slight effects on NK cell differentiation in vitro. This observation is consistent with previous studies of CD200-deficient mice, in which loss of CD200 predominantly altered myeloid cell activation and inflammatory responses without reported defects in immune-cell development [92]. Thus, like the activating receptor, NKp80, which does not have an established role in lymphopoiesis but serves as a useful marker to denote committed and functionally competent NKDIs [17, 18], CD200R1 appears to be useful in refining the human NK cell developmental pathway, serving as a marker that can distinguish uncommitted precursors from otherwise mature NK cells within the broader and previously described S4A population.

In summary, we identify CD200R1 as a previously unrecognized marker of human NKDIs that distinguishes immature, oligopotent S4A cells from a more mature CD200R1⁻ population with advanced transcriptional, DNA methylation, and functional NK cell features. While CD200R1 marks distinct developmental states and tissue communication niches, the receptor itself appears largely dispensable for NK cell differentiation. Together, these findings refine the current model of human NK cell development by revealing previously unrecognized heterogeneity within the S4A compartment and linking developmental state to lineage potential, epigenetic programming, and microenvironmental interactions.

## LIMITATIONS OF THE STUDY

First, our analyses were performed in the context of human tonsil tissue, which may not fully reflect NK cell developmental dynamics in other anatomical sites. Second, although our single-cell transcriptomic, DNA methylation, and ligand-receptor inference analyses identify associations between CD200R1 expression and distinct developmental and microenvironmental states, these approaches are correlative and do not establish direct causal relationships. Moreover, ligand-receptor and niche interaction analyses were computationally inferred and require experimental validation. Third, while in vitro differentiation and functional assays support developmental plasticity within the CD200R1⁺ compartment, these systems may not fully recapitulate the complexity of in vivo niche interactions.

## RESOURCE AVAILABILITY

### Lead Contact

Further information and requests for resources and reagents should be directed to and will be fulfilled by the lead contact, Aharon Freud.

### Materials Availability

All unique/stable reagents generated in this study are available from the lead contact with a completed materials transfer agreement.

## ACKNOWLEDGEMENTS

We thank Victoria Sellers and Lauren Smith for administrative and lab supervisory assistance. We are grateful to the donors who contributed deidentified tonsil samples to this study. This work was supported by grants from the National Institutes of Health (R01 CA208353 and R01 AI198432 to A.G.F., R01 CA255860 to B.L.M.-B. and C.C.O., and F30 CA236063 to A.P.N.) as well as funds from The Ohio State University Comprehensive Cancer Center (OSUCCC)-affiliated Pelotonia Organization (to A.P.N., M.R.L., B.L.M.-B., and A.G.F.) and the 2024 AACR-AstraZeneca Endometrial Cancer Research Fellowship (Grant Number 24-40-12-YOUS to Y.Y.). Y.Y. was also supported by the American Society of Hematology (ASH) Research Training Award for Fellows (RTAF). Research reported in this publication was also supported by The OSUCCC and the National Institutes of Health under award number P30 CA016058. We thank the staff of the Flow Cytometry Shared Resource at The OSU Comprehensive Cancer Center, which is supported by the NCI (P30CA016058).

## AUTHOR CONTRIBUTIONS

Y.Y., A.P.N., and M.A.R. contributed equally to this work. They conceived the study and led its design, performed experiments and data analyses, generated figures, and wrote the original draft of the manuscript. M.R.L., N.T.K. D.C., J.C., M.B., and E. A. performed experiments. W.C. and A.G.M provided experimental expertise and guidance and edited the final manuscript. X.Z. oversaw and performed statistical analyses. J.S.B. and B.W.B. facilitated scRNAseq data generation and analyses. C.C.O. oversaw the generation and analysis of DNA methylation assays. B.L.M.-B. provided expertise on NK cell and ILC developmental cultures, lentivirus generation, and infection. A.G.F. and B.K. directed the study, secured funding, and wrote the manuscript. All authors reviewed and approved the final manuscript.

## DECLARATION OF INTERESTS

The authors declare no competing interests.

### Declaration of generative AI and AI-assisted technologies in the manuscript preparation process

Prior to the initial submission and during the initial preparation of this work, the authors used Gemini (Google) to refine the clarity, flow, and grammatical precision of the manuscript text. After using this tool/service, all authors reviewed and edited the content as needed. All authors take full responsibility for the content of the published article.

## SUPPLEMENTAL INFORMATION

**Document S1: Figures S1 – S2.**

## STAR METHODS TEXT

### Human Tissue Samples

Human tissues were collected and used in accordance with protocols approved by The Ohio State University Institutional Review Board (OSU IRB). Human pediatric tonsils were obtained fresh following same-day surgeries via the NCI-approved Cooperative Human Tissue Network (CHTN) from Nationwide Children’s Hospital, Columbus, OH or from Vanderbilt University Medical Center, Nashville, TN. Leukocyte-depleted red blood cells and leukocyte reduction system apheresis cones were commercially obtained from the American Red Cross and Versiti.

### Cell Isolation and Fluorescence-Activated Cell Sorting

Human ILCs and their developmental intermediates were enriched from fresh tonsil tissue specimens using previously reported protocols [93]. Briefly, single cell suspensions from fresh human pediatric tonsils were generated by dissociation via a GentleMACS Dissociator (Miltenyi Biotec) according to manufacturer’s instructions. Cells were resuspended in fetal bovine serum (FBS, Sigma-Aldrich) and mixed with leukocyte-depleted red blood cells (RBCs) and a custom human NK cell enrichment RosetteSep reagent containing a cocktail of bivalent antibodies against glycophorin A and CD3, CD4, CD19, CD20, CD36, CD66b, and CD123 (STEMCELL Technologies). Cells mixed with RBCs and RosetteSep reagent were incubated on a nutator for 30 min at room temperature, diluted in PBS (Thermo Fisher Scientific), layered over Ficoll-Paque PLUS (GE Healthcare), and centrifuged at 1800 rpm for 30 min at room temperature with the brake off. The mononuclear cell layers were harvested and residual RBCs were lysed using eBioscience 1x RBC Lysis Buffer (Thermo Fisher Scientific). The tonsillar ILC-enriched fractions were then either used immediately for phenotypic and functional studies or individual NKDI subsets were sorted to >99% purity using a FACSAriaII cell sorter (BD Biosciences). S3 cells (aka ILCPs) were sorted as live, lineage (Lin) antigen (Lin = CD3, CD4, CD5, CD14, CD19, CD20, CD123, FcεR1α, TCRαβ, TCRδγ) negative, CD34⁻CD117^+^CD94⁻NKp80⁻CD16⁻ NKp44⁻CD294⁻KLRG1⁻ lymphocytes. S4A cells were sorted as live Lin⁻CD94^+^NKp80⁻CD16⁻ lymphocytes. S4B cells were sorted as live Lin⁻CD94^+^NKp80^+^CD16⁻ lymphocytes. Additionally, and when indicated in the text, CD200R1⁺ and CD200R1⁻ subsets of S3 and S4A NKDI populations were sorted.

### In Vitro Cell Cultures

All in vitro cell culture experiments were incubated at 37°C in 5% CO_2_ atmosphere. For bulk culture experiments, FACS-purified tonsil-derived ILCs were seeded at 1000 cells per well and cultured with IL-7 and IL-15 (10 ng/ml each; Miltenyi Biotec) for 28 days in 200 µl culture media per well in 96-well flat-bottom plates (TrueLine). For single cell clonal assays, FACS-purified populations were sorted directly into 96-well round-bottom plates, using established laboratory protocols for single cell clonal developmental assays [11, 12]. Briefly, cells were monitored via light microscope for visible identification of viable cell populations and subsequent harvest for analysis by flow cytometry. Cloning efficiency was approximately 10%, with a total of 49 clones analyzed from 4 donors across 2 independent experiments. Harvested clones with fewer than 10 live CD45^+^ events were excluded from final analyses. The culture media for in vitro development experiments contained DMEM and F12 (2:1 ratio) supplemented with 1% antibiotic/antimycotic (Thermo Fisher Scientific), 20 mg/ml ascorbic acid, 24 µM 2-mercaptoethanol, 0.05 mg/ml sodium selenite (Sigma), and 10% heat-inactivated human AB serum (Valley Biomedical). For experiments with stromal feeder cells, OP9-DL1 cells were originally obtained from the lab of Dr. Juan Carlos Zuniga-Pflucker [94]. Stromal cells were maintained in MEM-α ^+^ Glutamax media (Thermo Fisher Scientific) with 10% fetal bovine serum and 1% antibiotic/antimycotic (Thermo Fisher Scientific). One day prior to culture of NKDIs, OP9-DL1 cells were seeded at a density of 1000 cells per well and incubated at 37°C for cell adhesion. Stromal cell media was removed immediately prior to addition of FACS-sorted ILCs. Fresh media and cytokines were replenished twice per week. For intracellular cytokine analysis, cells were resuspended in 250 µl RPMI-1640 media and rested for 16 hours overnight at 37°C, after which they were stimulated for 4 hr with either IL-2 (10 ng/ml, PeproTech) plus a mixture of 81 nM phorbol 12-myristate 13-acetate (PMA) and 1.34 µM ionomycin (Cell Stimulation Cocktail, Thermo Fisher Scientific) for intracellular cytokine analysis, or cocultured with MHC class I–negative K562 target cells for NK cell degranulation analysis. A mixture of 10.6 µM brefeldin A and 2 µM monensin (Protein Transport Inhibitor Cocktail, Thermo Fisher Scientific) was also added during the 4 hr stimulation.

### Flow Cytometry

*Ex vivo* and in vitro-derived ILC populations were stained using antibodies directed against surface +/_⁻_ intracellular proteins according to manufacturer’s instructions. Where appropriate, isotype-matched or unstimulated controls were used to determine nonspecific staining. LIVE/DEAD Fixable Aqua Dead Cell Stain Kit and SYTOX Blue Dead Cell Stain (Thermo Fisher Scientific) viability dyes were used to exclude non-viable cells in the analyses. Intracellular staining was performed using Cytofix and Cytoperm Fixation and Permeabilization Solution Kit (BD Biosciences) for cytokine analysis, or Foxp3 Transcription Factor Staining Buffer Set (Thermo Fisher Scientific) for transcription factor analysis. Data were acquired on an LSRII cytometer (BD Biosciences) or Attune NXT (Thermo Fisher Scientific) and analyzed using FlowJo (BD Biosciences) software.

### Single Cell RNA-Sequencing, Library Generation, Sequencing, and Analysis

For Figure 1, single cell RNA-Sequencing data from enriched tonsil ILCs NKDIs was acquired from GEO dataset GSE242100 [26]. Fastq files were pre-processed using cellranger v6.0.2. Further analysis was performed in Seurat [95] with scCustomize for further streamlined analysis (10.5281/zenodo.5706430) [96]. Poor quality cells were defined as those with greater than three median absolute deviations above or below the median level in the following metrics: percent mitochondrial genes, percent genes in top 20 expressed genes for the entire dataset, number of counts per cell, number of genes per cell. Doublets were removed using scDblFinder [97]. PCA, clustering, and dimensional reduction were performed in Seurat. Trajectory analysis was performed using Monocle 3 [98]. Cell type was determined by mapping of gene modules based on differential gene expression analyses of bulk RNAseq datasets from the following studies: GSE169644 [46], GSE112813 [99], and GSE176367 [15]. Counts were normalized and lowly expressed genes were filtered out. Differential gene expression analysis was performed using the limma package [100]. The top 100 genes for each bulk sorted cell type were then mapped onto the tonsil scRNAseq data with the AddModuleScore function.

For Figure 2, the predicted S3 subset present in the original UMAP was isolated and re-clustered and further differentiated based on CD200R1 gene expression (mean expression >0.4 was positive).

For Figure 3, FACS-sorted S4A NKDIs from three donors were fixed in methanol and stored at −80 degrees Celsius prior to processing. Cells were washed and rehydrated in the presence of DTT and RNase inhibitor according to published protocols [101]. Single cells were isolated using the 10X Genomics 3’ v2 kit and a Chromium controller. Barcoded cDNA and libraries were generated according to the kit manufacturer’s instructions. Sequencing data were pre-processed using cellranger v3.1.0. Sample quality control was performed using *scater* [102].

The scRNAseq data were analyzed in Python using Scanpy [103]. Following quality control filtering and doublet removal, cells were normalized, log-transformed, and integrated across donors using Harmony [104]. Dimensional reduction was performed by principal component analysis, followed by neighborhood graph construction, Leiden clustering, and UMAP visualization. Only clusters 1, 2, and 3 were kept for downstream analysis due to poor quality of other cells present in the data. To infer transcription factor activity across S4A NKDIs, regulon activity was estimated using decoupleR [105] with the CollecTRI [106] transcription factor-target interaction database.

For Figure 6, cell-cell communication analyses were performed using LIANA[107]. S4A NKDIs were also analyzed together with previously annotated immune, stromal, and endothelial cell populations from human tonsillar single-cell datasets from the following studies: human tonsil atlas (https://doi.org/10.5281/zenodo.10373041) [55], endothelial cell atlas (GSE124494) [56]. Ligand-receptor interactions were inferred using the LIANA rank-aggregation framework, and interactions predicted to target S4A NK cell subsets were compared between CD200R1-expressing and CD200R1-negative populations. Differential ligand-receptor interactions, pathway enrichment analyses, and predicted cellular sources of signaling were subsequently used to define microenvironmental niches associated with CD200R1^+^ and CD200R1^-^ S4A subsets.

### Lentiviral Production and Overexpression of CD200R1 Ligand (CD200) in OP9-DL1 Stromal Cells

HEK293T cells were used for lentiviral production and were cultured in Dulbecco’s Modified Eagle Medium (DMEM) supplemented with 10% Fetal Bovine Serum (FBS) and antibiotic/antimycotic (Thermo Fisher Scientific). All cells were maintained in a humidified incubator at 37°C with 5% CO2. Twenty-four hours before transfection, 5×10^6^ HEK293T cells were seeded in a T-75 flask to achieve 70-80% confluency. For viral packaging, HEK293T cells were co-transfected with a 4:2:1 molar ratio of the transfer plasmid (Control: pLV[Exp]-EGFP/Neo-EF1A; CD200: Control Vector + hCD200; VectorBuilder), the packaging plasmid (psPAX2; Addgene #12260), and the envelope plasmid (pMD2.G; Addgene #12259), respectively. A total of 20 µg of plasmid DNA was transfected using Lipofectamine 3000 (Invitrogen #L3000015) following the manufacturer’s protocol. The transfection medium was replaced with 10 mL of fresh growth medium 12-16 hours post-transfection. Viral supernatant was collected at 48 and 72 hours post-transfection, pooled, and centrifuged at 500 x g for 10 minutes to pellet cellular debris. The clarified supernatant was then filtered through a 0.45 µm sterile PES syringe filter. Filtered virus was concentrated 10-fold with PEG-it (Systems Bioscience) according to the manufacturer’s instructions. Pellets were resuspended in cold PBS and stored at −80°C. Cell titer was determined by transduction of HEK293T cells with serially diluted virus and then detection of GFP via flow cytometry. OP9-DL1 cells were infected with control or CD200 overexpression virus using a Multiplicity of Infection (MOI) of 10 with 8 µg/mL of polybrene. Virally transduced cells were chemically selected via culture with 100 µg/mL geneticin for 1 week, followed by FACS sorting of OP9-DL1 cells expressing CD200 to achieve a pure population.

### Methylation Array

DNA was extracted from freshly sorted tonsil NKDI subsets using the Gentra Puregene Cell Kit (Qiagen). Genome-wide DNA methylation was generated using Infinium MethylationEPIC v1.0 Kit (Illumina) according to manufacturer’s recommended procedures. Data were processed using the RnBeads software [108] and normalized using BMIQ [109]. Illumina probes located on X/Y chromosomes, non-CpG probes, and probes with a SNP (>0.01 MAF) located ≤5 bp to the analyzed CpGs were censored for a total of 804,572 analyzed CpGs. Principle component analysis (PCA) was performed on the top 5000 most variable probes from S3, CD200R1^+^ S4A, CD200R1⁻ S4A, and S5 NKDIs from tonsil. De novo motif enrichment from hypo and hyper methylated regions was determined using HOMER [110]. CpGs of interest were compared with a background probe set composed of probes matched for CpG, GC, and DNA methylation content; universe appropriateness was evaluated using the LOLA R package [111].

### Data and Code Availability

Single cell RNA sequencing (scRNA-seq) analysis scripts are available on our github repository found at https://github.com/blaserlab/freud_nk. New single cell data generated for this paper will be accessible through the NIH GEO following peer-review and subsequent publication.

