## Supplemental Figures for "CD200R1 Marks a Branch Point During Human Natural Killer Cell Development in Tonsils"

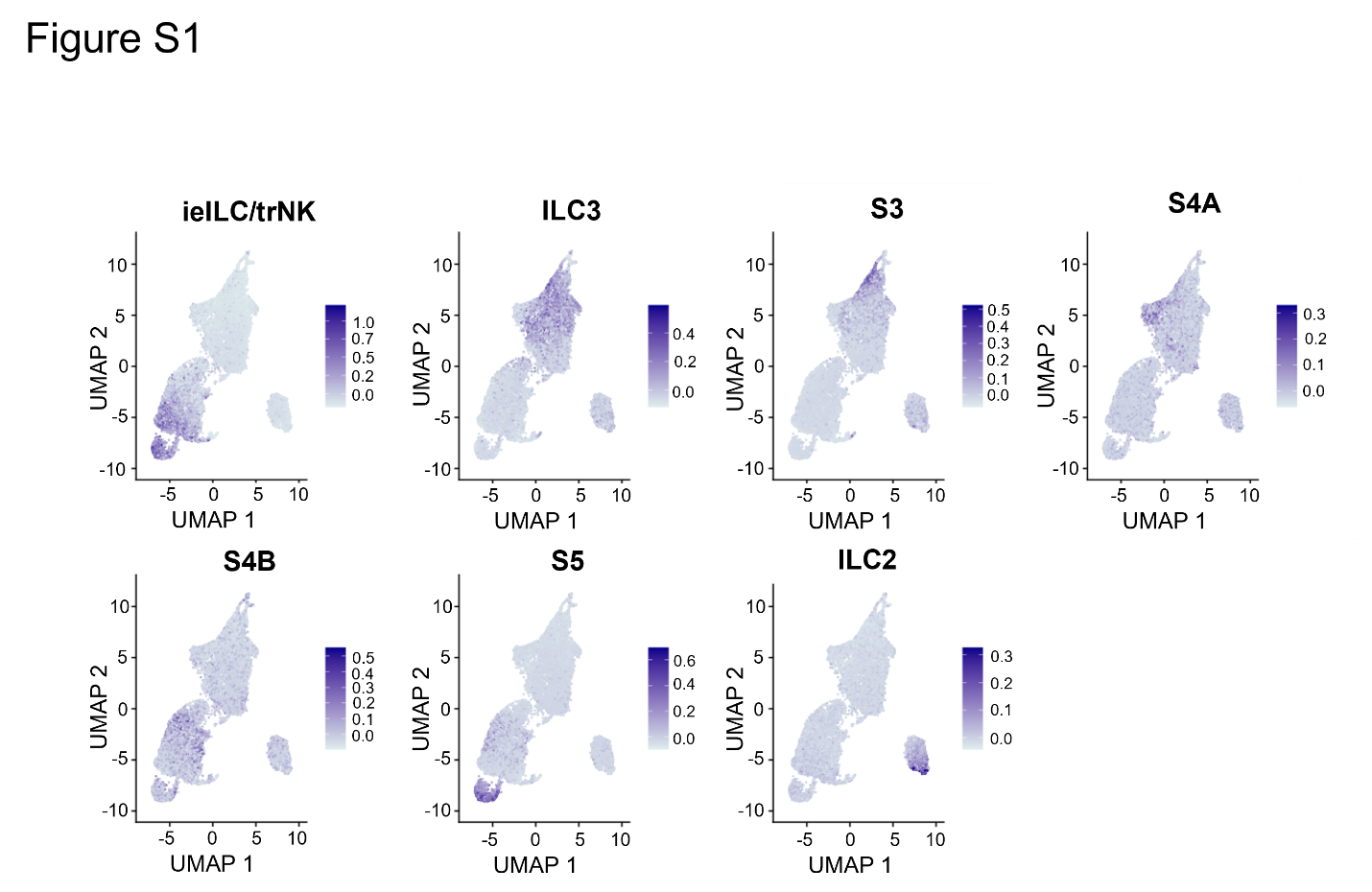

**Figure S1.** **Bulk RNA-seq-derived gene expression modules mapped to corresponding NKDI and ILC populations in tonsil scRNA-seq data.**

UMAP feature plots depicting module scores generated from publicly available bulk RNA-seq datasets for the indicated NKDI and ILC populations. Differentially expressed genes (DEGs) for each population were identified using limma-voom, and module scores generated from the top upregulated 200 DEGs for each population were mapped onto the tonsil scRNA-seq dataset shown in Figure 1A. See also Figure 1.

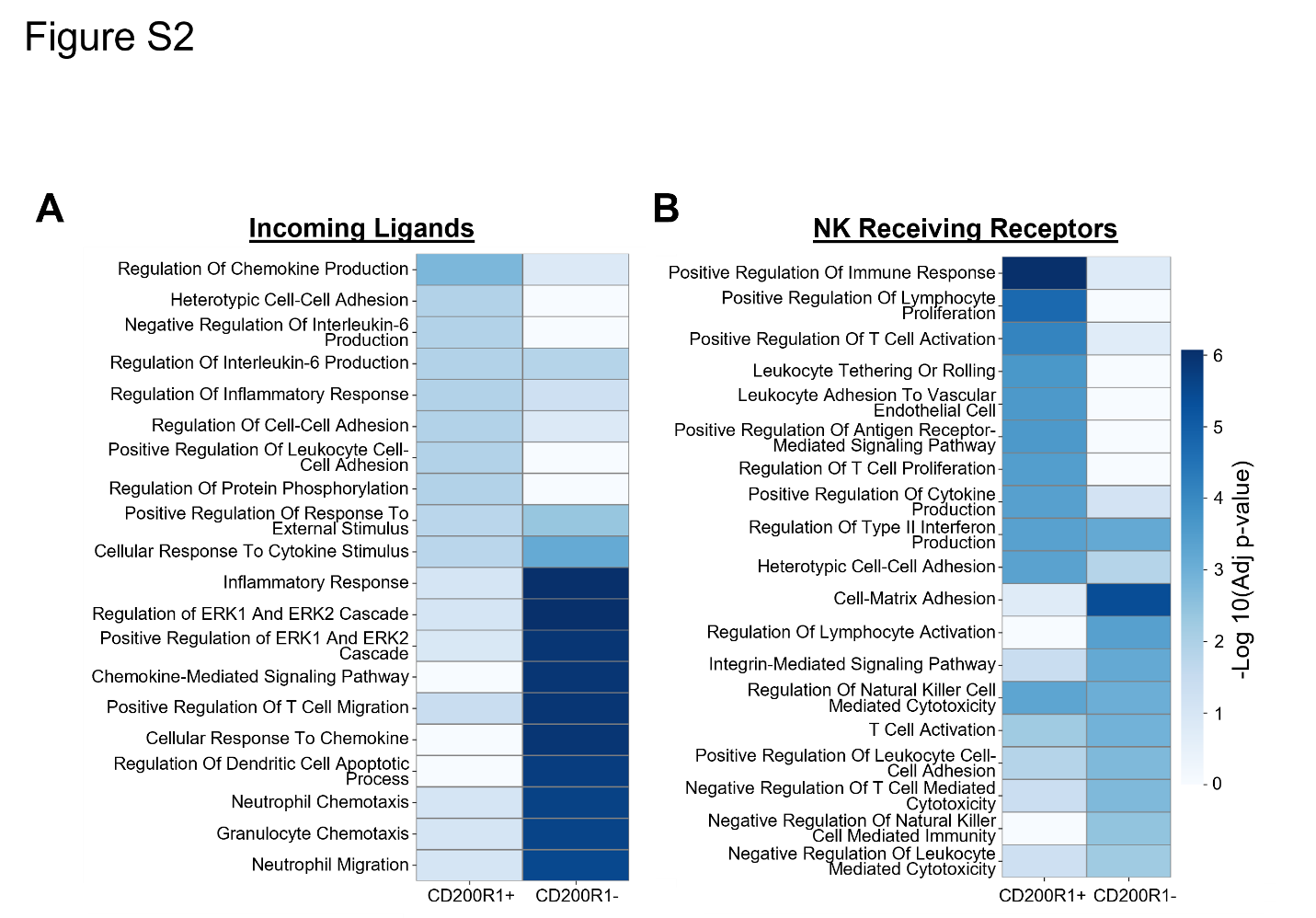

**Figure S2**. **Gene Ontology (GO) enrichment analysis of scRNA-seq-derived predicted niche interactions involving CD200R1⁺ and CD200R1⁻ S4A NKDIs.**

(A) GO Biological Process enrichment analysis of incoming ligands associated with CD200R1⁺ or CD200R1⁻ S4A niche interactions. Color intensity represents −log₁₀(adjusted P value).

(B) GO Biological Process enrichment analysis of NK cell receiving receptors associated with CD200R1⁺ or CD200R1⁻ S4A niche interactions. Color intensity represents −log₁₀(adjusted P value).
